# Autoreactive IgG levels and Fc receptor γ subunit upregulation drive mechanical allodynia after nerve constriction or crush injury

**DOI:** 10.1101/2025.03.22.644748

**Authors:** Nathan T. Fiore, Kendal F. Willcox, Anamaria R. Grieco, Dorsa Dayani, Younus A. Zuberi, Cobi J. Heijnen, Peter M. Grace

## Abstract

B cells contribute to the development of pain after sciatic nerve chronic constriction injury (CCI) via binding of immunoglobulin G (IgG) to Fc gamma receptors (FcγRs) in the lumbar dorsal root ganglia (DRG) and spinal cord. Yet the contribution of B cells to pain after different types of peripheral nerve injury is uncertain. Using male and female mice, we demonstrate a divergent role for B cell-IgG-FcγR signaling underlying mechanical allodynia between CCI, nerve crush (NC), spared nerve injury (SNI), and spinal nerve ligation (SNL). Depletion (monoclonal anti-CD20) or genetic deletion (muMT mice) of B cells prevented development of allodynia following NC and CCI, but not SNI or SNL. In apparent contradiction, circulating levels of autoreactive IgG and circulating immune complexes were increased in all models, though more prominent following NC and CCI. Passive transfer of IgG from SNI donor mice induced allodynia in CCI muMT recipient mice, demonstrating that IgG secreted after SNI is pronociceptive. To investigate why pronociceptive IgG did not contribute to mechanical allodynia after SNI, we evaluated levels of the Fc receptor γ subunit. SNI or SNL did not increase γ subunit levels in the DRG and spinal cord, whereas CCI and NC did, in agreement with B cell-dependent allodynia in these models. Together, the results suggest that traumatic peripheral nerve injury drives secretion of autoreactive IgG from B cells. However, levels of cognate FcγRs are increased following sciatic nerve constriction and crush, but not transection, to differentially regulate pain through the B cell-IgG-FcγR axis.

## 1. Introduction

Neuropathic pain is a debilitating condition that arises from lesion or disease affecting the somatosensory system [35; 60]. Its treatment is challenging due to the diverse underlying causes and mechanisms driving pain development [12]. Thus, there is a need for expanded understanding of the underlying mechanistic heterogeneity and related therapeutic targets [19; 57]. The development of neuropathic pain following nerve injury is driven by a complex network of interactions among neurons, glia, and immune cells [20; 27; 39]. Accumulating preclinical and clinical evidence indicates that (auto)antibodies produced by B cells can promote nociceptive signaling via Fc gamma receptor (FcγR) signaling, complement activation, or disruption of ion channels expressed by DRG neurons, thereby driving neuropathic pain [14; 28; 45; 46; 53; 58]. We recently reported that the depletion of B cells (via anti-CD20 administration or genetic deletion) prevented mice from developing mechanical allodynia, highlighting a pro-nociceptive role for B cells in the CCI model [46]. In contrast, B cells are reported to be unnecessary for the development of mechanical allodynia after spared nerve injury (SNI) or spinal nerve ligation (SNL), since mice lacking mature B cells, or nude mice (lacking both T and B cells) reconstituted with T cells, develop mechanical allodynia after SNI or SNL injury respectively [8; 13]. Therefore, it remains unclear whether B cells contribute to the development of allodynia specifically following CCI, or in various other models of peripheral nerve injury as well.

In mice, CCI induces secretion of immunoglobulin G (IgG) which engage functional Fc gamma receptors (FcγRs), which are required for the development of mechanical allodynia after CCI [46]. Importantly, we also found that IgG accumulates in the DRGs from human donors with chronic pain, as well as in the DRGs and spinal cord following CCI in mice [46]. IgG accumulation has also been reported at the site of injury following sciatic nerve crush (NC) in mice and rats [71; 77]. We also reported that CCI increases expression of the Fc receptor γ subunit (FcRγ), a key component of FcγRs, in the DRG and spinal cord [21]. Similarly, NC increases cellular expression of FcRγ and FcγRs at the injured nerve and DRG, though no change in FcγR expression has been reported in the DRG following SNI or SNL [4; 33; 38; 59; 73; 78; 79]. Given that FcγR signaling primarily occurs from engagement with immune complexes (ICs), the pronociceptive effects of IgG are likely mediated through formation of complexes with antigen [6; 45]. However, the identity of antigens released, the binding affinity of secreted IgG to autoantigens, and formation of ICs after nerve injury remains to be determined. Here, we aim to characterize the role of B cell-IgG-FcγR signaling in various peripheral nerve injury models, specifically nerve crush (NC), compression (CCI), and ligation and transection (SNI and SNL).

## 2. Methods

### 2.1. Animals

Male and female C57BL/6J (Jackson Laboratory, strain #000664) or muMT mice (Jackson Laboratory, strain B6.129S2-*Ighm^tm1Cgn^*/J, strain #002288) were used for all experiments. For experiments with mice lacking mature B cells, muMT mice that have homozygous mutations in the immunoglobulin mu heavy chain (*Ighm^tm1Cgn^*) that leads to an absence of mature B cells were used [41]. MuMT mice were bred in-house at the University of Texas MD Anderson Cancer Center AAALAC accredited vivarium and wild-type (WT) littermate mice were used as controls. Mice were housed in the pathogen-free, MD Anderson vivarium, with *ad libitum* access to food and drinking water on a 12 h light-dark cycle. Mice were at least 8 weeks old at the start of experimental manipulations. Animals were prospectively assigned to treatment groups using an online group randomization tool (randomlists.com/team-generator). All animal experiments and procedures were approved by the Institutional Animal Care and Use Committee of the University of Texas MD Anderson Cancer Center.

### 2.2. B cell depletion via anti-CD20

Anti-CD20 (mouse IgG2a clone 5D2, Genentech CA) monoclonal antibodies (mAb) or isotype control antibody (IgG2a, Genentech) was delivered to mice via intravenous (i.v.) tail vein injections, as previously described (200 µg in 100 µL sterile 0.9% saline) [46]. Injections were delivered within 1 hr before NC, SNI or SNL surgeries.

### 2.3. Peripheral nerve injury models

Neuropathic pain was modelled using unilateral sciatic nerve CCI, NC, SNI or SNL as previously described [3; 5; 15; 21; 32; 40]. Mice were anesthetized with inhaled isoflurane (2.5% in 2L/min oxygen for induction and maintenance) and placed on an electric heating pad to maintain body temperature. The left hindlimb was shaved and cleaned with povidone-iodine and 70% ethanol. An incision was made to expose the left sciatic nerve and polypropylene pipette tips were used to isolate the sciatic nerve proximally to the nerve trifurcation. For sham surgeries, the sciatic nerve was isolated, but no nerve ligation or crush were applied. The muscle layer was sutured close (4-0 silk; Ethicon), and the skin was closed with a 9 mm wound clip. At the completion of surgery, mice were returned to their home cage and monitored postoperatively until fully ambulatory.

#### 2.3.1. Chronic constriction injury

For CCI surgeries, three ligatures (5-0 chromic gut; Ethicon, NJ USA) were loosely tied around the sciatic nerve approximately 1mm apart [5; 21; 46].

#### 2.3.2. Sciatic Nerve Crush

For NC surgeries, the full width of the sciatic nerve was crushed for 15 s using a modified ultra-fine hemostat (53096 Ted Pella, CA USA) with double aluminum foil layers forming a 30 μm spacer surrounding the nerve, placed on the first lock position [40].

#### 2.3.3. Spared Nerve Injury

For SNI surgeries, the tibial and common peroneal nerves were isolated, tightly ligated (6-0 silk, Ethicon), and transected, with the sural nerve left intact [3; 15].

#### 2.3.4. Spinal Nerve Ligation

For SNL surgeries, the L5 transverse process was removed to expose L4 and L5 spinal nerves. The L5 spinal nerve was isolated, tightly ligated (6-0 silk, Ethicon) and transected [32].

### 2.4. Behavioral tests for mechanical allodynia

Animals were habituated to the testing apparatus and the experimenter prior to experimentation. The experimenter was blinded to group assignment of each animal throughout testing. Von Frey testing of punctate allodynia was assessed using the up-down method, and 50% withdrawal thresholds were calculated, as previously described [9; 21]. After von Frey tests were complete, dynamic allodynia was measured by lightly stroking the plantar surface of the hindpaw with a soft paintbrush (5/0, Princeton Art & Brush Co., WI USA), as previously described [21]. Average scores for each mouse were obtained from three stimulations at intervals of at least 15 s.

### 2.5. Serum autoantigen microarray

Blood was collected via cardiac puncture following a lethal injection of pentobarbital-phenytoin solution (i.p.) 14 days post-surgery. Blood was centrifuged at 3200 x g at room temperature for 10 mins, and serum was collected and stored at -80°C. 10μL of serum was used to profile IgG autoantibodies using the Autoantigen Microarray Super Panel (128 antigens). Serum was treated with DNAse I, diluted 1:50, and incubated with the autoantigen array. The autoantibodies binding to the antigens on the array were detected with Cy3 labelled anti-IgG and the arrays scanned with GenePix® 4400A Microarray Scanner. The images were analyzed using GenePix 7.0 software to generate GPR files. The averaged net fluorescent intensity of each autoantigen was normalized to internal controls (IgG) to produce a normalized signal intensity (NSI), as well as the signal to noise ratio (SNR) for all antigens. Antigens with SNR < 5 were removed prior to analysis. Analysis was performed on NSI values, and Z scores was calculated by subtracting the sample NSI value from mean NSI and dividing by standard deviation for all sham, CCI, NC and SNI samples.

### 2.6. ELISAs

Blood was collected via cardiac puncture following a lethal injection of pentobarbital-phenytoin solution (i.p.) 14 days post-surgery, and serum isolated as described above. Following blood collection, mice were transcardially perfused with 10mL of ice-cold 0.9% saline and left L4/5 DRGs and left lumbar spinal cords were isolated, frozen in liquid nitrogen, and stored at -80°C. Serum concentration of soluble ICs was assessed using a circulating IC ELISA (EM2596-6 Lifeome, TX USA). Circulating IC ELISA was performed following manufacturer’s instructions, with serum diluted 1:50 and 100μL of diluted serum loaded per sample. All samples and standards were run in duplicate and IC concentration quantified based on the standard curve. IC concentration was expressed as ng per μL serum sample. To validate that the IC ELISA used was sensitive to serum concentrations of IgG-ICs, we diluted 100 μL of sham and CCI serum 1:1 with 8% polyethylene glycol-8000 (PEG8000, PO131-500GM Spectrum Chemical and Laboratory Products, CA USA) to reach a final PEG8000 concentration of 4%. 200 μL mixtures were vortexed and incubated overnight at 4 °C. The resulting precipitate was centrifuged at 2000 x g for 20 mins at 4 °C. The supernatant was collected for analysis, and the pellets washed with 200 μL of 4% PEG8000 solution, then centrifuged at 2000 x g for 20 mins at 4 °C. Supernatant was discarded and the pellet dissolved in 100 μL of sterile 1x phosphate buffered saline (PBS, pH 7.4, 10010-023 Thermo Fisher Scientific, MA USA). Circulating IC ELISA was performed as described above, with the collected supernatant or precipitate diluted 1:50 and 100μL of diluted supernatant or precipitate loaded per sample. Total IgG levels in the left L4/5 DRGs and left lumbar spinal cords were analyzed using a mouse IgG uncoated ELISA kit (88-50400-77 Invitrogen, MA USA) as previously described [21; 46]. Protein was isolated via chemical homogenization in 200 μL RIPA buffer (89900 Thermo Fisher Scientific) containing 1x Halt protease inhibitor cocktail (78430 Thermo Fisher Scientific) and mechanically homogenized using tissue pestles followed by 3 x 30 s cycles on tissue Bioruptor (Diagenode, NJ USA). Total protein was quantified via Pierce BCA protein assay kit (23225 Thermo Fisher Scientific). IgG ELISAs were performed following manufacturer’s instructions, with all samples and standards run in duplicate, and IgG concentration quantified relative to the total protein loaded for each sample (ng/pg).

### 2.7. IgG purification

IgG was purified from serum collected from WT mice on day 14 after CCI or SNI surgery as previously described [46]. Due to the large number of mice required to generate donor IgG, these studies were restricted to a single sex (male), and serum was pooled across subjects from the same surgical conditional. IgG was purified from serum using MabSelect SuRe resin (17543801 Cytiva, MA USA), with 5 mL of diluted serum (1:20 in sterile 1x PBS, pH 8) loaded onto the column containing 1 mL of the resin and elution performed with 0.1 M glycine and 0.1 M CaCl2 (pH 3). Protein concentration of collected elution was read using a Nanodrop (Thermo Fisher Scientific) at 280 nm, and the IgG samples were concentrated to 2 mg/mL via Amicon Ultra-4 10K (UFC801008 Millipore Sigma, MA USA), sterilized by filtration through 0.2 µm filter and then stored at 4°C.

### 2.8. Passive transfer of IgG

IgG was delivered to mice via intrathecal (i.t.) injections as previously described [46]. Male mice were used for these experiments, to match the sex of the IgG donors. Mice were anesthetized with inhaled isoflurane, placed on an electric heating pad to maintain body temperature, shaved along the lumbar region and skin cleansed with 70% ethanol. CCI IgG or sham IgG (40 µg in sterile PBS) was slowly injected (injection time ∼2 mins) into the intrathecal space above the lumbar enlargement using custom injectors (27G needle fitted to polyethylene tubing, PE-20, attached to 25 µL Hamilton syringe). MuMT recipient mice received 3x daily IgG injections beginning 14 days post-CCI. Pre von Frey measurements were taken 24 hrs before the first injection (day 13 post-CCI) and Post measurements taken 24 hrs after the final injection (day 17 post-CCI).

### 2.9. Immunohistochemistry

Mice were euthanized with an injection of pentobarbital-phenytoin solution (i.p.) 14 days post-injury and transcardially perfused with 10 mL of ice-cold 0.9% saline, followed by 10 mL of 4% paraformaldehyde (41678-0030 Thermo Fisher Scientific) in 0.1 M phosphate buffer (PB, pH 7.4). L4/5 DRG and lumbar spinal cords were post-fixed in 4% paraformaldehyde overnight prior to being cryoprotected stepwise in 15%, 22.5% and 30% sucrose in 0.1 M PB with 0.01% sodium azide (014314-09 Thermo Fisher Scientific) at 4°C. Tissues were freeze-mounted in O.C.T compound (4583 Sakura Finetek, CA USA), and serially sectioned (12μm for DRGs and 16μm for spinal cords) using a cryostat (Leica Biosystems, Nussloch Germany). Tissue sections were mounted to SuperFrost Plus charged slides (22-037-246Thermo Fisher Scientific) and stored at -20°C. For DRG, slides were washed in 0.3% Triton X-100 (T9284 Millipore Sigma) in 1x PBS (PBST) three times and incubated with blocking buffer (5% normal donkey serum (ab7475 Abcam, Cambridge UK), in PBST) for 1 hrs prior to primary antibody staining. For spinal cord, sections were permeabilized in 0.1% NaBH_4_ (452882 Millipore Sigma) in PBS for 20mins and washed twice (PBST) prior to blocking. Primary antibodies were added after blocking and incubated at 4°C overnight. Sections were then washed three times, incubated with secondary antibodies for 2 hrs, washed three times, incubated with DAPI (1 μg/mL; D9542 Millipore Sigma) for 5mins, washed in PBS four times, and cover slipped with FluorSave mounting medium (345789, Millipore Sigma). Primary antibodies used were goat anti-mouse IgG antibody (1:100; Jackson ImmunoResearch, PA USA), mouse anti-FcRγ (1:200, MBL Life science, Tokyo Japan) and rabbit anti-MAP2 (1:250, Abcam). Secondary antibodies used were donkey anti-goat antibody, donkey anti-mouse and donkey anti-rabbit antibody (1:500; Thermo Fisher Scientific). A list of primary antibodies and secondary antibodies is provided in **Table S1**.

### 2.10. Confocal imaging and analysis

To assess IgG and FcRγ expression in the ipsilateral L4/5 DRG and spinal cord dorsal horn, fluorescent photomicrographs were captured using a Nikon Eclipse Ti2 Confocal Microscope (Nikon, Tokyo Japan) at 20X magnification for DRGs or 40X magnification for spinal cords. Z-stacks were acquired for each tissue section and maximum intensity projections of IgG were analyzed using ImageJ v1.53t software. Regions of interest were manually drawn around the DRG (excluding nerve root) or left dorsal horn spinal cord, and the mean gray intensity of IgG and FcRγ measured. Two to three sections were analyzed per animal, and the average mean gray intensity calculated for each animal.

### 2.11. Quantitative real-time PCR

To determine expression levels of *Fcer1g*, encoding the Fc receptor γ subunit, mice were transcardially perfused with 10mL of ice-cold 0.9% saline following a lethal injection of pentobarbital-phenytoin solution 7 days post-injury. Left L4/5 DRGs and left dorsal lumbar spinal cords were isolated, frozen in liquid nitrogen, and stored at -80°C. Total RNA from left L4/5 DRGs and left L4/5 spinal cords was isolated using RNeasy plus mini kit (74134 Qiagen, Hilden Germany) and cDNA was generated by reverse transcription using iScript gDNA Clear kits (1725035 Bio-Rad, CA USA) following manufacturer instructions. RT-qPCR was performed by diluting cDNA and associated primers in SsoAdvanced Universal SYBR Green Supermix (1725274 Bio-Rad) and monitoring the reaction with the CFX Connect Real-Time PCR Detection System (Bio-Rad). Relative gene expression against *Gapdh* using the 2^-ΔΔCT^ method was performed as previously described [22]. A list of primers is provided in **Table S1**.

### 2.13. Statistics

Data are presented as group mean ± SEM with individual subjects presented as symbols whenever feasible, unless otherwise indicated. Area under curve (AUC) (arbitrary units, a.u.) data are presented as box and whisker plots indicating median, inter-quartile range, and min/max. There were no sex differences, so data from both sexes were combined to form a single group. Statistical analyses were performed with GraphPad Prism software (v10.2.0). Von Frey data was analyzed by two-way repeated measures ANOVA and Tukey’s post hoc tests. The AUCs from Brush test data were analyzed by Mann-Whitney test. Immunofluorescence, qPCR, IgG and IC ELISA data were analyzed by one-way ANOVA and Dunnett’s post hoc test. Serum autoantigen microarray data were analyzed by using Significance Analysis of Microarrays (SAM, v5.0) in R (v4.3.1) [67]. Autoantigen NSI from CCI, NC or SNI IgG was compared to Sham IgG using SAM two-class unpaired two-tailed T test design with Benjamani-Hochberg correction to account for false discovery rate. Significant autoantigens were identified where fold change ≥ |2| and *q* < 0.05 relative to sham. MuMT FcRγ immunofluorescence data were analyzed by unpaired two-tailed T test. For all data points, circle shapes represent male (●) and triangles represent female (▲) mice. Differences between groups were considered statistically significant when *p* < 0.05, or *q* < 0.05 for autoantigen microarray data.

## 3. Results

### 3.1. B cells are required for mechanical allodynia after sciatic nerve crush but not spared nerve injury or spinal nerve ligation

We recently showed that B cells drive mechanical pain following sciatic nerve CCI in mice [46]. To determine whether B cells influence the development of allodynia in other peripheral nerve injury models, we depleted B cells at the time of NC, SNI, or SNL via a single intravenous injection of anti-CD20. A single anti-CD20 injection prevented the development of hindpaw punctate and dynamic allodynia for at least 49 days after NC injury (**Figure 1A-C**). As pharmacological agents can have off-target effects, we used muMT mice as a complementary genetic approach to further investigate whether B cells influence mechanical allodynia. After sciatic NC, neither punctate nor dynamic allodynia developed in the muMT mice compared with WT littermate controls (**Figure 1D-F**) confirming that B cells contribute to the development of mechanical allodynia after NC. In contrast, SNI- and SNL-injured mice treated with anti-CD20 developed mechanical allodynia, which persisted for at least 49 days after injury (SNI **Figure 2A-C** and SNL **Figure 2D-F**). Furthermore, SNI- and SNL-injured muMT mice also developed mechanical allodynia (SNI **Figure 2G-I** and SNL **Figure 2J-L**). These data demonstrate that B cells are required to induce mechanical allodynia after sciatic nerve crush, but not ligation and transection.

**Figure 1.**
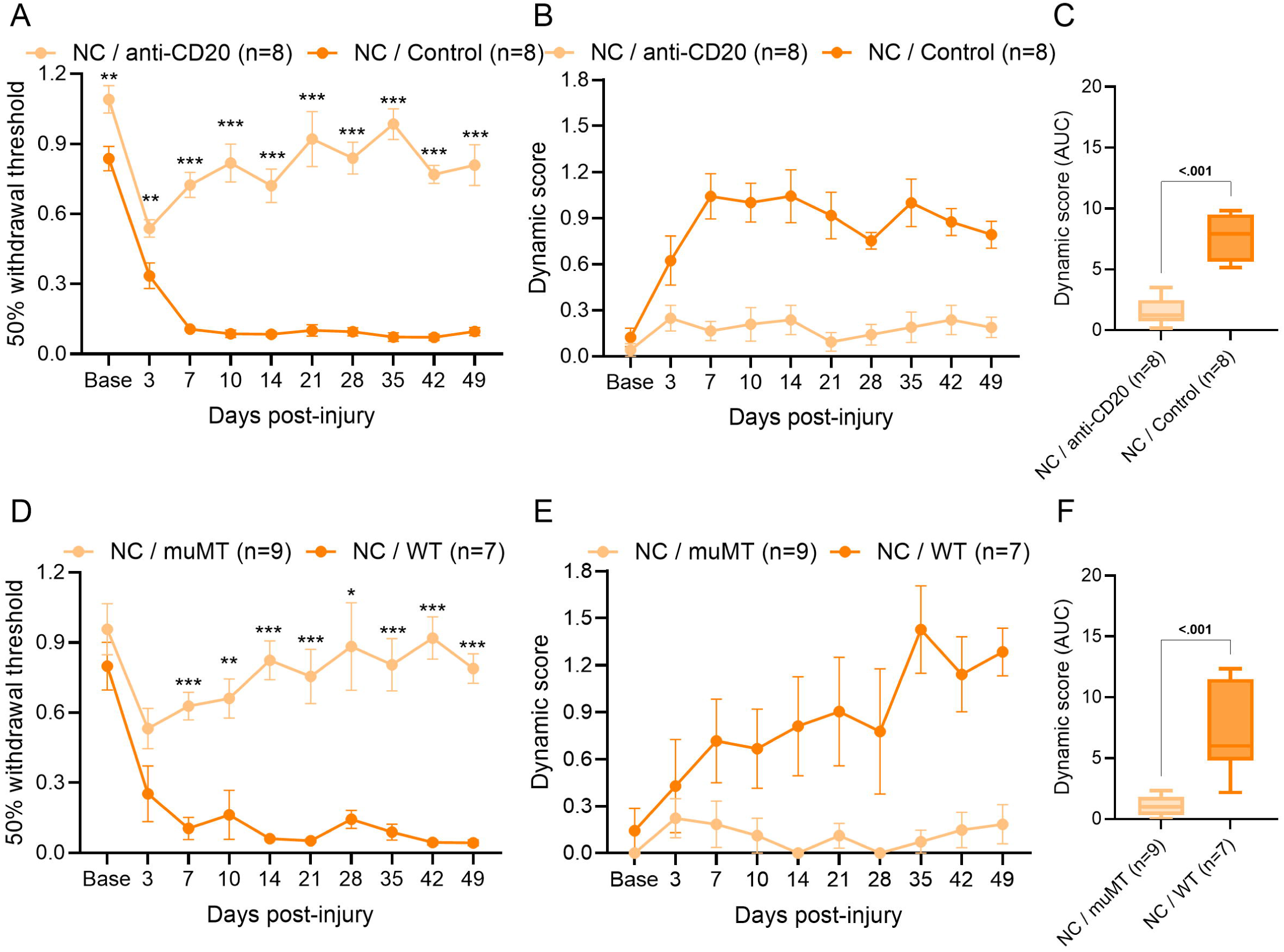
B cells are required for the development of mechanical allodynia after sciatic nerve crush. (**A**) Von Frey thresholds for punctate allodynia and (**B**) dynamic scores from brush test were assessed for ipsilateral hindpaws following i.v. anti-CD20 or IgG2a control (200μg/100μL) given at the time of nerve crush (NC). Von Frey analyzed by two-way repeated measures ANOVA and Sidak’s post hoc test; \**p* < 0.05, \*\**p* < 0.01, \*\*\**p* < 0.001 NC / anti-CD20 compared to NC / Control. *n* = 8/group (4 males and 4 females). (**C**) AUC of dynamic score (Baseline- Day 49) analyzed by Mann-Whitney test; *p* value shown. (**D**) Von Frey thresholds for punctate allodynia and (**E**) dynamic scores from brush test were assessed for ipsilateral hindpaws following NC in muMT or WT littermates. Von Frey analyzed by two-way repeated measures ANOVA and Sidak’s post hoc test; \**p* < 0.05, \*\**p* < 0.01, \*\*\**p* < 0.001 NC / muMT compared to NC / WT. *n* = 7-9/group (3-4 males and 4-5 females). (**F**) AUC of dynamic score (Baseline- Day 49) analyzed by Mann-Whitney test; *p* value shown.

**Figure 2.**
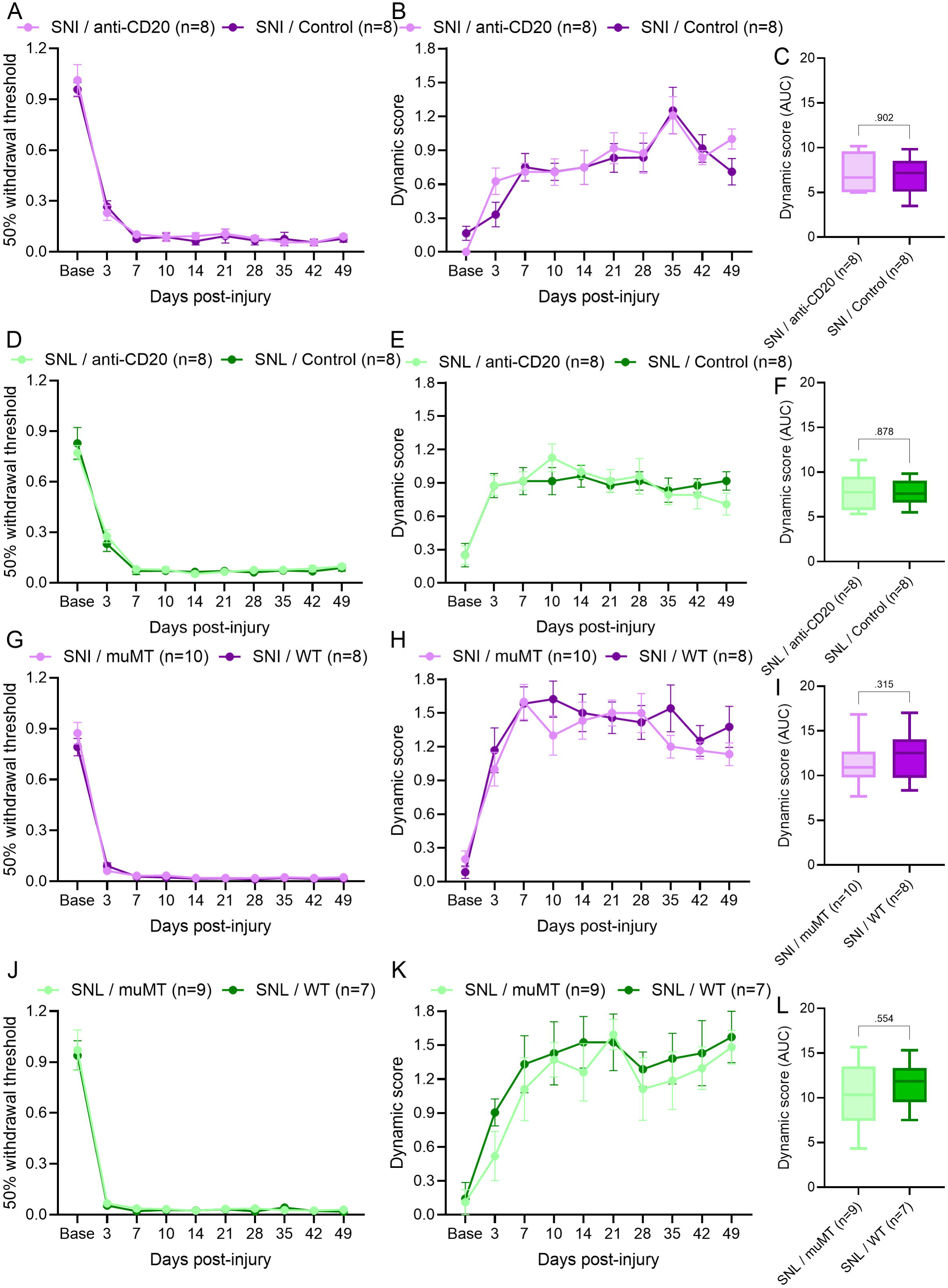
B cells are not required for mechanical allodynia after spared nerve injury or spinal nerve ligation. (**A**) Von Frey thresholds for punctate allodynia and (**B**) dynamic scores from brush test were assessed for ipsilateral hindpaws following i.v. anti-CD20 or IgG2a control (200μg/100μL) given at the time of SNI. *n* = 8/group (4 males and 4 females). (**C**) AUC of dynamic score (Day Base-49) analyzed by Mann-Whitney test; *p* value shown. (**D**) Von Frey thresholds for punctate allodynia and (**E**) dynamic scores from brush test were assessed for ipsilateral hindpaws following SNI in muMT or WT littermates. *n* = 8-10/group (4 males and 4-6 females). (**F**) AUC of dynamic score (Day Base-49) analyzed by Mann-Whitney test; *p* value shown. (**G**) Von Frey thresholds for punctate allodynia and (**H**) dynamic scores from brush test were assessed for ipsilateral hindpaws following i.v. anti-CD20 or IgG2a control (200μg/100μL) given at the time of SNL. *n* = 8/group (4 males and 4 females). (**I**) AUC of dynamic score (Day Base-49) analyzed by Mann-Whitney test; *p* value shown. (**J**) Von Frey thresholds for punctate allodynia and (**E**) dynamic scores from brush test were assessed for ipsilateral hindpaws following SNL in muMT or WT littermates. *n* = 8-10/group (4 males and 3-5 females). (**K**) AUC of dynamic score (Day Base-49) analyzed by Mann-Whitney test; *p* value shown.

### 3.2. Traumatic nerve injuries increase circulating levels of autoreactive IgG and soluble immune complexes

Given the differential regulation of injury-induced allodynia by B cells, we hypothesized that CCI and NC, but not SNI, would lead to increased levels of autoreactive IgG. SNI was selected as the representative ligation and transection model because it involves partial denervation of the sciatic nerve and is more comparable to CCI, NC and the sham injury control. In contrast, SNL introduces additional damage by requiring the removal of the L5 transverse process to isolate the spinal nerves. An autoantigen microarray panel was used to profile serum IgG reactivity against a broad range of autoantigens 14 days post-injury (**Figure 3A** and **Table S2**). We identified increased IgG reactivity towards 30 autoantigens after CCI, 30 after NC, and 12 after SNI compared to sham IgG (**Figure 3B** and **Table S3**). There was also decreased reactivity for 2 autoantigens after SNI relative to sham IgG (**Table S3**). Among the autoantigens with increased IgG reactivity, 6 were common to all injury groups, 20 were shared between CCI and NC, 3 were shared between NC and SNI, and 1 was shared between CCI and SNI (**Figure 3B**). The reactive autoantigens common to all injury groups included CD4, complement C4, neuropilin-1 (NRP1), ribosomal P0 protein (P0), E2 component of pyruvate dehydrogenase complex (PDC-E2) and proteinase 3 (PR3) (**Figure 3B** and **Table S3**).

**Figure 3.**
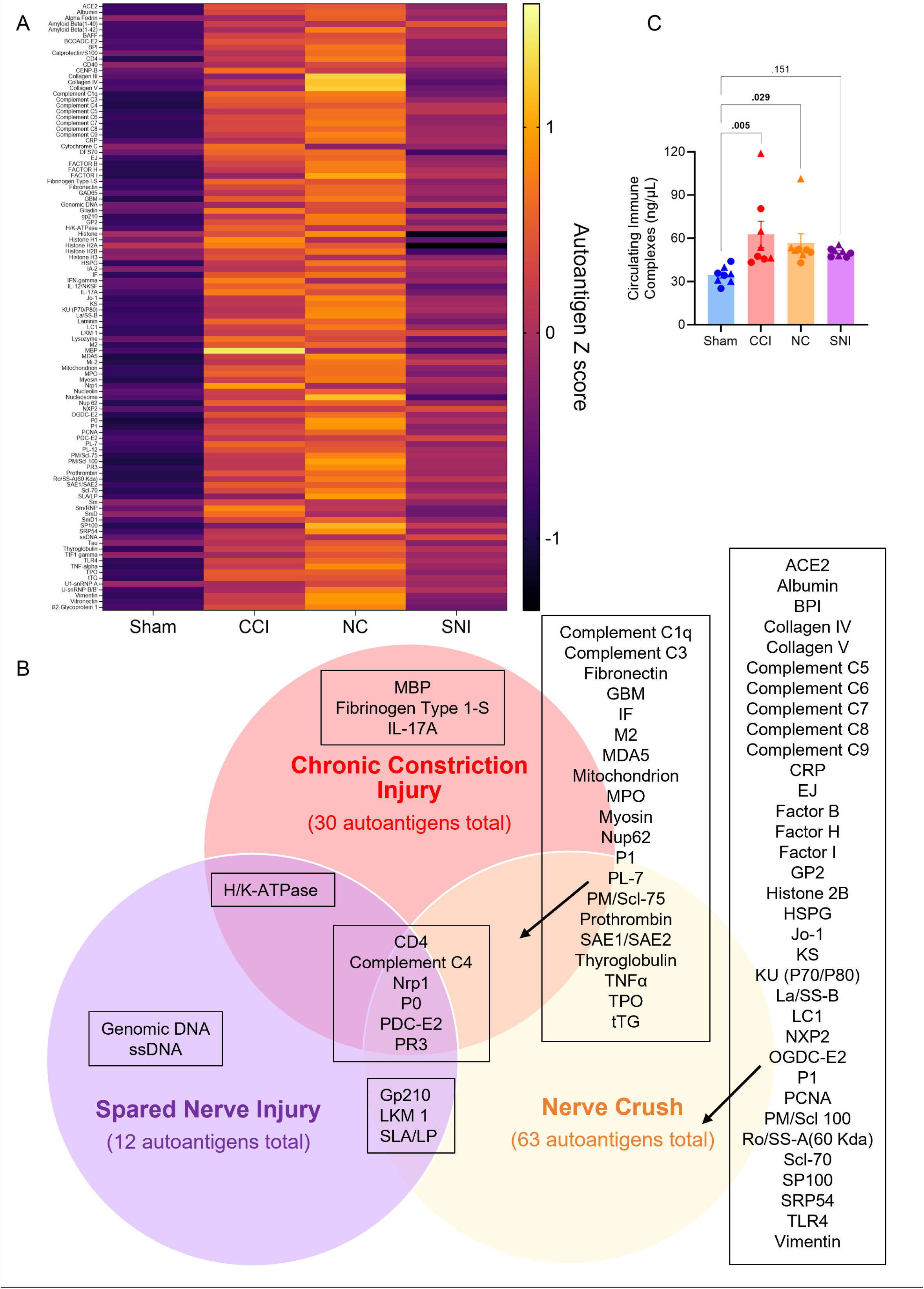
Traumatic nerve injury broadly increases levels of autoreactive IgG. (A) Z scores for serum IgG autoantigen reactivity 14 days after Sham, CCI, NC or SNI injury. (B) Venn diagram comparing autoantigens with increased IgG autoreactivity after CCI, NC and SNI. Autoantigen IgG reactivity was identified as upregulated relative to Sham using SAM two-class unpaired analysis (Fold change ≥ 2 and *q* < 0.05). *n* = 15/group (8 males and 7 females). (C) Quantification of circulating ICs protein levels in serum via ELISA. ELISA analyzed by one-way ANOVA and Dunnett’s post hoc test; *p* values shown for comparisons to Sham. *n* = 8/group (4 males and 4 females). For all data points, circles (●) represent male and triangles (▴) represent female mice.

To determine whether the autoreactive IgG led to formation of ICs, we assessed circulating ICs 14 days post-injury. We observed an increase in IC levels in the serum after CCI and NC compared to sham-injured mice (**Figure 3C**). There was a modest non-significant increase after SNI (**Figure 3C**), which correlated with the low reactivity towards autoantigens (**Figure 3A-B**). To validate our results, we analyzed ICs after precipitating out the IgG complexes from serum with 4% PEG8000 [2; 10; 52]. We confirmed that the concentration of CCI ICs was reduced in the PEG8000 supernatant, compared to the isolated PEG8000 precipitate (**Figure S1**). Collectively, these data highlight that all nerve injury models result in the secretion of autoreactive IgG, contrasting with our original hypothesis. However, IgG secreted after CCI and NC is reactive towards a greater number of autoantigens, and has higher affinity towards those antigens, than that secreted after SNI. Consistently, a greater number of ICs are formed after CCI and NC compared to SNI.

### 3.3. IgG secreted after SNI is pronociceptive

Since SNI did result in secretion of autoreactive IgG, albeit to a lesser extent than other models, we tested its pronociceptive capacity. We purified IgG from the serum of CCI or SNI donor mice (WT) (referred to as CCI-IgG or SNI-IgG). MuMT recipient mice underwent CCI, followed by passive transfer of SNI-IgG or CCI-IgG (40 μg i.t. per day on days 14, 15, and 16 post CCI). SNI-IgG induced punctate allodynia in recipient muMT mice, relative to pre-injection thresholds (**Figure 4A**). The positive control CCI-IgG also induced punctate allodynia, consistent with prior observations (**Figure 4B**) [46]. These data indicate that IgG secreted after SNI is pronociceptive.

**Figure 4.**
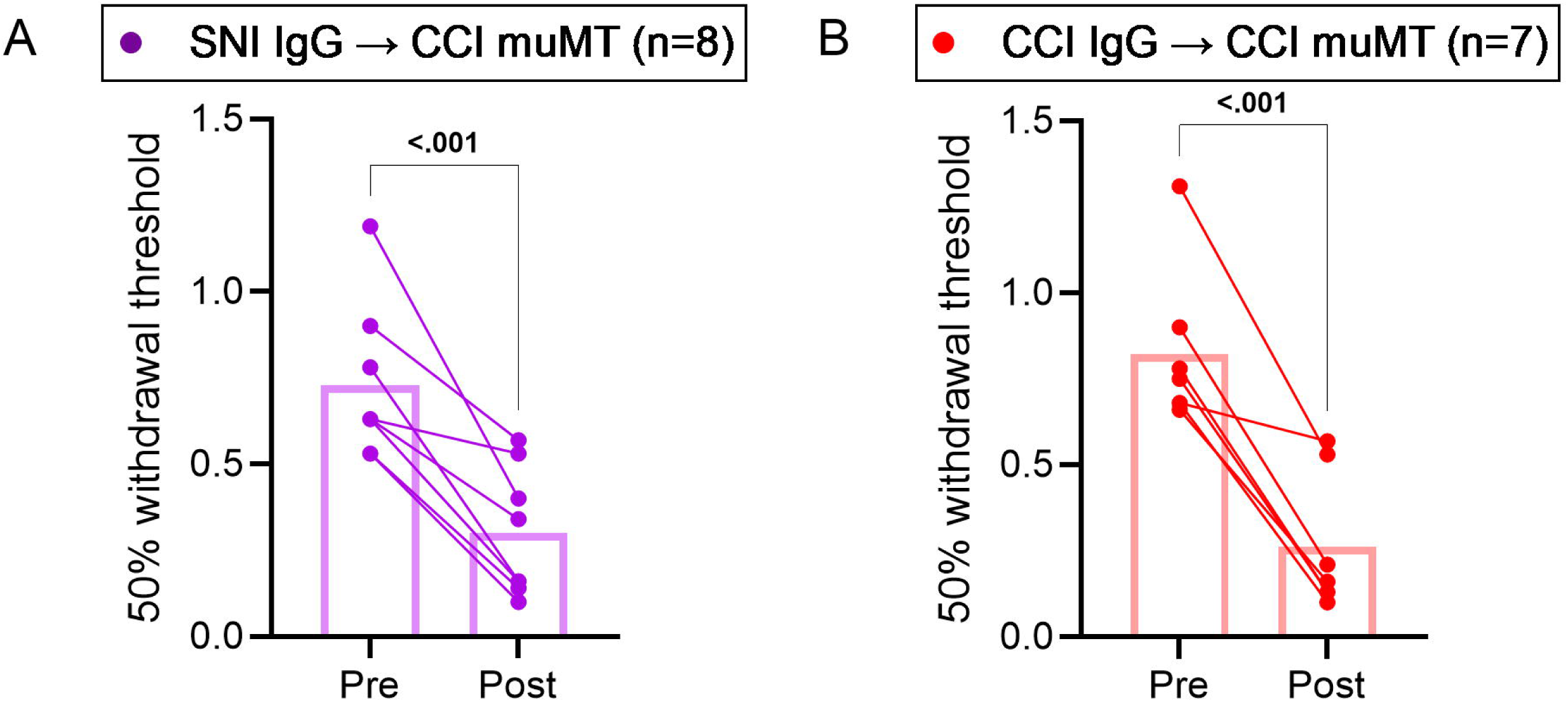
IgG secreted after SNI is pronociceptive. Passive transfer of CCI or SNI IgG induces allodynia in injured recipient mice. IgG from WT CCI or SNI donor mice were passively transferred to CCI-muMT recipient mice (3 x 40 μg daily i.t. injections starting 14 days post-CCI). Von frey thresholds for punctate allodynia were assessed for ipsilateral hindpaws before (Pre) and 24 hrs after the last injection (Post). Von Frey analyzed by two-way repeated measures ANOVA and Sidak’s post hoc test; *p* value shown for Pre compared to Post. *n* = 7-8/group (7-8 males).

### 3.4. IgG accumulates in the ipsilateral DRG and spinal cord after CCI and NC but not after SNI or SNL

We have previously shown that IgG accumulates in the DRG and spinal cord after CCI which likely drives mechanical pain based on the sufficiency of intrathecal IgG transfer to induce allodynia into injured muMT mice (**Figure 4B**) [46]. To understand whether IgG accumulation around the lumbar region is correlated with the differential role of B cells across nerve injury models, we measured IgG levels in the ipsilateral L4/5 DRG and spinal cord dorsal horn 14 days after injury using immunohistochemistry and ELISA. IgG immunoreactivity and IgG protein levels were increased in the ipsilateral DRG (**Figure 5A-C**) and spinal cord dorsal horn (**Figure 6A-C**) after CCI and NC, relative to sham controls. Following SNI and SNL, IgG immunoreactivity was only modestly increased in the ipsilateral DRG relative to sham controls (**Figure 5A-B**). However, ELISA measurements showed that IgG protein levels remained unaltered after SNL and SNI (**Figure 5C**). Similarly, IgG immunoreactivity was moderately increased in the spinal cord dorsal horn after SNL relative to sham controls but remained unaltered after SNI (**Figure 6A-B**). Additionally, IgG protein levels in the spinal cord were unchanged for both SNI and SNL (**Figure 6C**). These data suggest that CCI and NC lead to greater IgG accumulation in the DRG and spinal cord than SNI or SNL.

**Figure 5.**
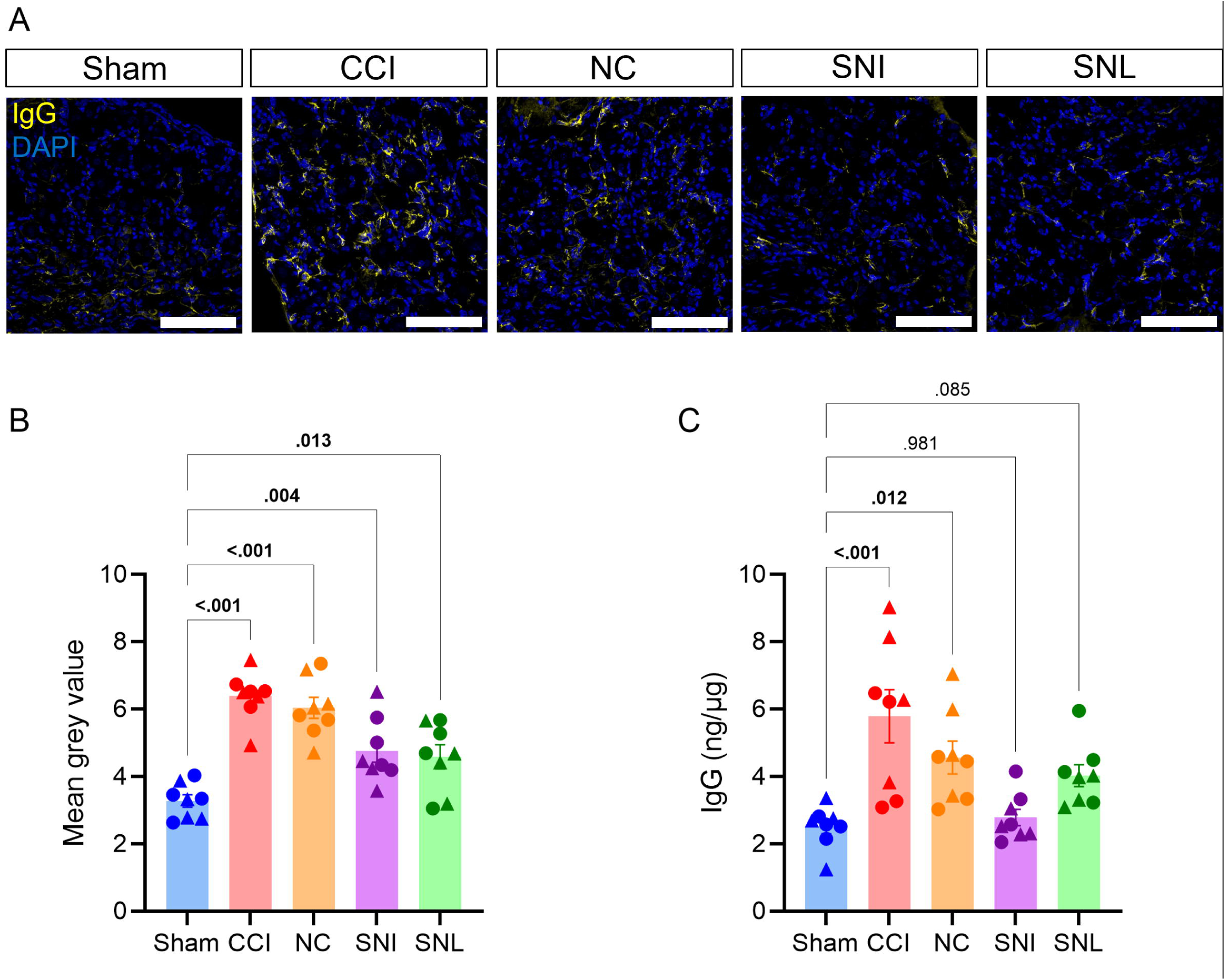
IgG accumulation increases in ipsilateral DRG after CCI and NC, but not after SNI or SNL. IgG accumulation in the ipsilateral lumbar DRG after sham, CCI, NC, SNI or SNL. (**A**) Representative 40x fluorescent images of IgG (yellow) and DAPI (blue) in the ipsilateral lumbar DRG (male, day 14 after surgery). Scale bars indicate 100μm. (**B)** Quantification of mean gray intensity for IgG. (**C**) Quantification of IgG protein levels in DRG via ELISA. One-way ANOVA and Dunnett’s post hoc test; *p* values shown for comparisons to Sham. *n* = 8/group (4 males and 4 females). For all data points, circles (●) represent male and triangles (▴) represent female mice.

**Figure 6.**
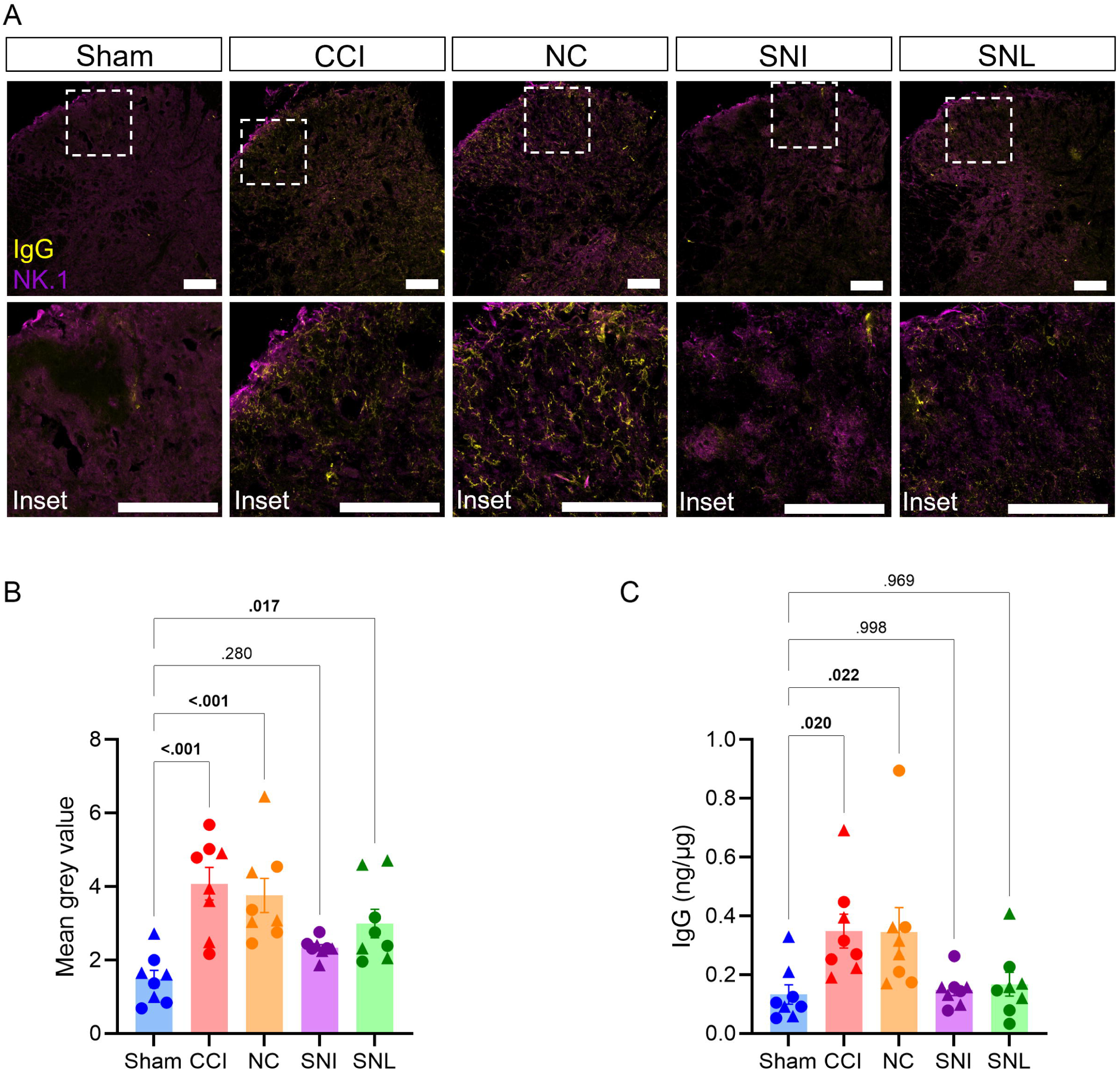
IgG accumulation increases in ipsilateral dorsal horn after CCI and NC, but not after SNI or SNL. IgG accumulation in the ipsilateral lumbar spinal cord dorsal horn after sham, CCI, NC, SNI or SNL. (**A**) Representative IgG (yellow) and NK.1 (magenta) 20x fluorescent images of ipsilateral spinal cord dorsal horn with 60x inset image (male, day 14 after injury). (**B)** Quantification of mean gray intensity for IgG. (**C**) Quantification of IgG protein levels in lumbar spinal cord via ELISA. One-way ANOVA and Dunnett’s post hoc test; *p* values shown for comparisons to Sham. *n* = 8/group (4 males and 4 females). For all data points, circles (●) represent male and triangles (▴) represent female mice.

### 3.5. CCI and NC but not SNI or SNL, upregulate FcR**γ** subunit levels in the ipsilateral DRG and spinal cord

The differential accumulation of IgG in DRG and spinal cord across nerve injury models may be due to varying FcγRs levels. We therefore assessed protein immunoreactivity and gene expression of Fc γ subunit (FcRγ), which is common to all FcγRs that contain an immunoreceptor tyrosine-based activation motif (ITAM) in our four models of nerve injury [54]. In the ipsilateral DRG, FcRγ protein immunoreactivity was increased 14 days after CCI and NC, compared to sham (**Figure 7A-B**). FcRγ gene (*Fcer1g*) expression was also increased in DRG at 7 days after CCI and NC, compared to sham (**Figure 7C**). There were similar trends in the spinal cord dorsal horn for FcRγ immunoreactivity (**Figure 7D-E**) and *Fcer1g* expression (**Figure 7F**). SNI and SNL did not lead to an increase in FcRγ immunoreactivity or gene expression in the DRG and spinal cord, compared to sham controls (**Figure 7A-F**). Primary antibody controls were used in DRG sections to confirm the specificity of IgG (**Figure S2A**) and FcRγ (**Figure S2B**) immunolabeling in WT mice. These data suggest that CCI and NC, but not SNI or SNL, upregulate FcRγ gene expression and protein immunoreactivity in the ipsilateral DRG and spinal cord at 7- and 14-days post-injury, respectively.

**Figure 7.**
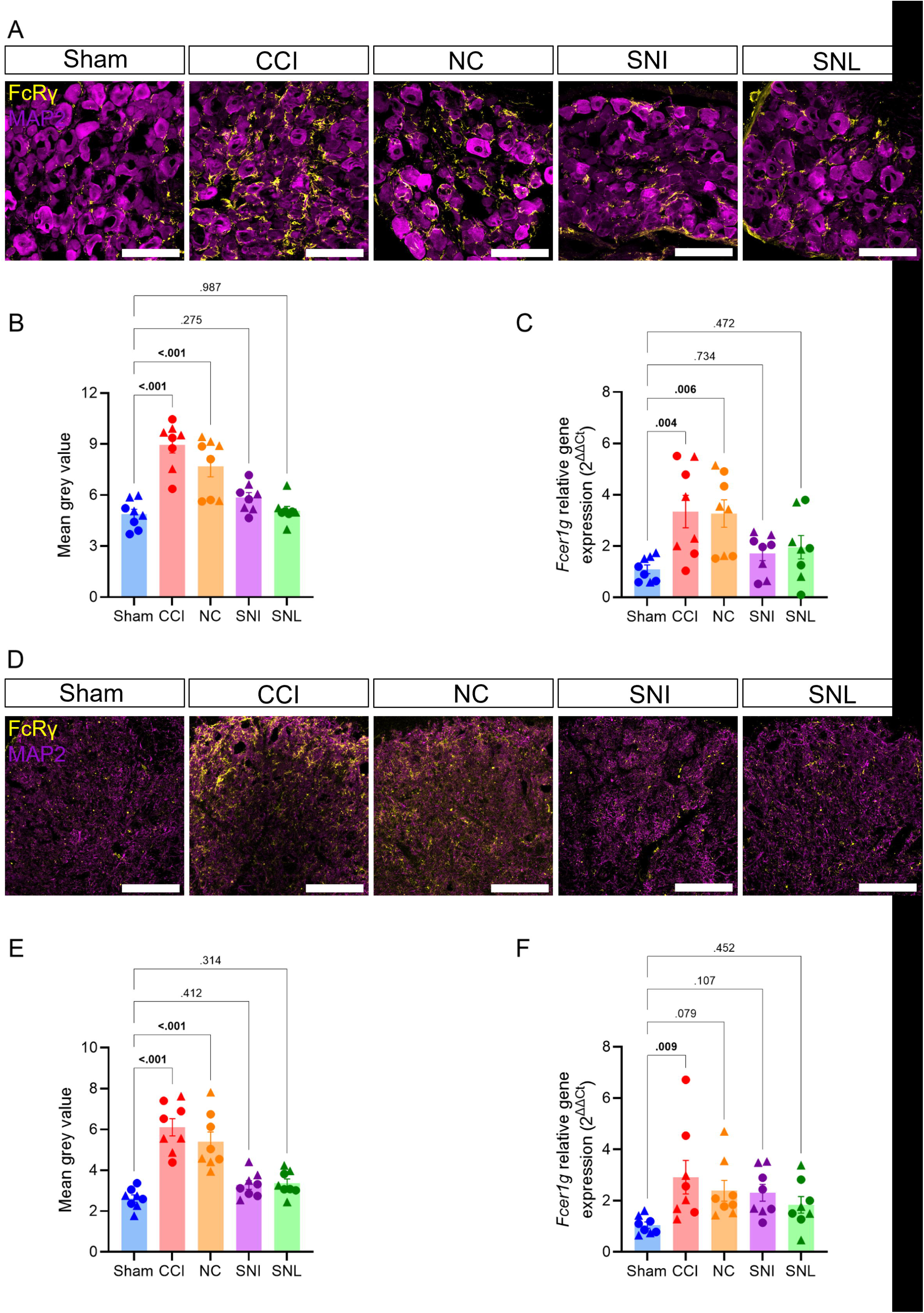
Only CCI and NC upregulate Fc_γ_ subunit in ipsilateral DRG and spinal cord. FcRγ accumulation in the ipsilateral lumbar DRG and spinal cord dorsal horn after sham, CCI, NC, SNI or SNL. (**A**) Representative 40x fluorescent images of IgG (yellow) and MAP2 (magenta) in the ipsilateral lumbar DRG (male, day 14 after surgery). Scale bars indicate 100μm. (**B)** Quantification of mean gray intensity for FcRγ. (**C**) Quantification of relative FcRγ (*Fcer1g*) gene expression against Gapdh using the 2^-ΔΔCT^ method in ipsilateral lumbar DRG. (**D**) Representative 40x fluorescent images of IgG (yellow) and MAP2 (magenta) in the ipsilateral spinal cord dorsal horn (male, day 14 after surgery). Scale bars indicate 100μm. (**B)** Quantification of mean gray intensity for IgG. (**C**) Quantification of relative FcRγ (*Fcer1g*) gene expression against Gapdh using using the 2^-ΔΔCT^ method in ipsilateral spinal cord dorsal horn 7 days post-injury. One-way ANOVA and Dunnett’s post hoc test; *p* values shown for comparisons to Sham. *n* = 8/group (4 males and 4 females). For all data points, circles (●) represent male and triangles (▴) represent female mice.

## 4. Discussion

We have identified that B cells differentially regulate neuropathic pain based on the nature of traumatic nerve injury. Cellular depletion or genetic deficiency of B cells protected mice from developing mechanical allodynia after NC, complementing our previous findings with CCI [46]. In contrast, these interventions did not modify the trajectory of mechanical allodynia after SNI or SNL. CCI and NC increase circulating levels of IgG with increased reactivity towards 20 common autoantigens. These findings were accompanied by an increase in circulating IgG immune complexes (ICs). In apparent contradiction to the behavioral results, SNI still led to an increase in circulating autoreactive IgG. Passive transfer experiments demonstrated that this IgG was pronociceptive. However, where IgG accumulated in the ipsilateral lumbar DRG and spinal cords after CCI or NC, there was no increase after SNI or SNL. In alignment with these data, levels of the Fc receptor γ subunit were increased after CCI and NC, but not SNI or SNL. Based on these results, we propose that traumatic peripheral nerve injury broadly induces secretion of autoreactive IgG. However, levels of FcγRs are increased following constriction and crush but not transection, which may represent the point of divergence in the pronociceptive B cell-IgG-FcγR axis for different traumatic nerve injuries.

Traumatic nerve injury broadly led to increased serum IgG reactivity towards various autoantigens. This increased IgG autoreactivity was accompanied by an increase in circulating ICs after CCI and NC, highlighting that nerve injury induces the production of autoreactive IgG which form antigen-antibody ICs. Elevated concentrations of circulating ICs have been linked to disease progression in various autoimmune conditions, including rheumatoid arthritis and systemic lupus erythematosus [1; 48; 65]. Autoreactive IgG contributes to the development of pain in various autoimmune conditions, including complex regional pain syndrome, fibromyalgia, rheumatoid arthritis and collagen antibody-induced arthritis [18; 24; 31; 37; 44; 49; 64]. Since mice lacking the Fc γ chain subunit do not develop mechanical allodynia following passive transfer of IgG from CCI donors, autoreactive IgG secreted following nerve injury likely mediates pain via FcγR-dependent signaling [46]. The production of anti-myelin antibodies has been reported following CCI or NC [11; 47; 71]. Here we identified increased reactivity to MBP exclusively in CCI IgG. Given that IgG from SNI donors mediates mechanical allodynia when passively transferred to muMT recipients after CCI, increased IgG reactivity towards common antigens likely drives these nociceptive effects. Autoantigens with increased IgG reactivity across all nerve injury models include intracellular (such as the mitochondrial protein PDC-E2, PR3 in neutrophil cytoplasm, and ribosomal protein P0 in the nucleus), and extracellular antigens (such as complement C4, CD4 on helper T cells, and Nrp1, which is expressed by neurons, blood vessels, and various immune cells). Autoantibodies against these self-antigens have previously been reported in autoimmune conditions, such as systemic lupus erythematosus, where autoreactive IgG contributes to disease progression and symptoms, including pain [25]. The de novo presence of intracellular self-antigens as targets for production of autoantibodies are probably derived from apoptotic or necrotic cells which release intracellular contents into the extracellular environment via extracellular vesicles, resulting in formation of apoptotic bodies or extracellular traps [62]. Peripheral nerve injury likely leads to excessive release of apoptotic bodies or extracellular traps due to insufficient clearance, allowing these self-antigens to persist in the extracellular environment [42; 62; 66]. This sustained presence enables their recognition by B cells, ultimately driving autoantibody production [62].

Furthermore, the process of apoptosis induces several posttranslational modifications in the nucleus, including histone citrullination and acetylation, which can promote cross-reactivity between intracellular and extracellular antigens [17]. Posttranslational modifications, such as citrulline-modified peptides, also generate extracellular neoantigens, leading to recognition by B cells and the production of autoantibodies that drive disease progression of rheumatoid arthritis [68]. Notably, modified antibodies associated with rheumatoid arthritis, in which citrulline has replaced arginine, recognize a variety of proteins that were also identified after nerve injury such as collagen, fibronectin, histones and vimentin [56; 69; 74]. Posttranslational modifications are also linked to nerve regeneration [51; 61]. Considering that crush injury leads to faster and more complete nerve regeneration compared to transection injury, increased extracellular protein modifications could underlie the differences in IgG autoreactivity between NC, CCI and SNI after injury[72]. Given the large number and variety of autoantigens identified, nerve injury likely generates autoreactive IgG that either recognizes shared antigenic epitopes or reacts to multiple distinct epitopes, contributing to neuropathic pain development following compression or crush injury.

The levels of FcRγ were elevated in the ipsilateral DRG and spinal cord after CCI and NC, likely leading to greater IgG accumulation at these sites. Our results align with preclinical evidence suggesting that FcγRs in the DRG and spinal cord are more upregulated in CCI or NC models compared to transection injury models [4; 16; 22; 26; 33; 36; 38; 46; 50; 59; 63; 73; 78; 79]. FcRγ is a subunit of FcγRI, FcγRIII, and FcγRIV, all of which activate intracellular signaling cascades via ITAMs and spleen tyrosine kinase (Syk) [54]. The clustering of FcγRs triggers phosphorylation of their ITAM domain and activation of Syk family kinases, leading to the induction of proinflammatory signaling pathways [6]. In the context of nerve injury, Syk activation in the DRG and spinal cord is critical to the development of mechanical allodynia, as evidenced by Syk inhibitors dampening downstream inflammatory signaling and alleviating pain following both CCI and SNI [23; 50]. FcRγ expression fluctuates in response to injury and subsequent inflammatory signaling [7]. Given that SNI or SNL does not alter FcRγ expression, and that CCI- or SNI-derived IgG re-establishes pain in muMT mice after CCI, it is likely that IgG aggregate accumulation and/or differential release of proinflammatory chemokines and cytokines (such as interferon-γ) in the DRG and spinal cord after CCI or NC enhance the surface expression of ITAM-containing FcγRs [16; 21; 34; 55; 70]. However, a comprehensive comparison is necessary to identify the specific variations in the inflammatory milieu within the DRG and spinal cord that drive differential FcRγ expression after injury. Given that SNI-induced mechanical allodynia is Syk-dependent, it is likely that alternative upstream mechanisms leading to Syk activation compensate for lack of IgG-FcγR signaling to drive development of neuropathic pain following transection injury [23]. For example, colony stimulating factor 1 (CSF1) can mediate DNAX activating protein of 12kDa (DAP12) activation, which also contains an ITAM domain that activates Syk family kinases [30]. Given that axotomy results in the production of neuronal CSF1 and that DAP12 is required for the development of mechanical allodynia after SNI and SNL, it is evident that SNI and SNL lead to the production of neuronal CSF1 which engages DAP12 in DRG macrophages and spinal microglia to drive neuropathic pain [29; 43; 75; 76]. Therefore, while these different peripheral nerve injury models induce pain via Syk-dependent pathways, they likely activate Syk family kinases through distinct upstream mechanisms.

The extent to which neuropathic pain caused by various nerve injuries can be treated by targeting B cells remains an open question. Here, we demonstrate that cellular depletion (monoclonal anti-CD20) or genetic deficiency (muMT mice) of B cells prevented the development of allodynia following NC, but not SNI or SNL. Our results after NC align with our prior finding that B cells mediate pain after CCI, whereas the lack of effect following SNI or SNL aligns with reports that muMT mice develop mechanical allodynia after SNI, and that T cell reconstitution is sufficient to restore mechanical allodynia in nude mice after SNL [8; 13]. The enduring effect of a single injection of anti-CD20 mAbs at the time of CCI or NC, along with its lack of effect after SNI and SNL, suggests that targeting B cell-IgG-FcγR signaling may only alleviate pain following nerve constriction or crush injuries. Given our data showing no specific autoantigen and that B cell depletion does not reverse established mechanical allodynia after CCI, therapeutic strategies targeting pan-IgG clearance are preferable to B cell depletion or targeting a specific autoantibody. Such approaches are likely promising for alleviating pain following CCI or NC [46]. In support of this, we recently reported that reducing IgG accumulation in the DRG and spinal cord after CCI via blockade of the neonatal Fc receptor (FcRn) alleviates pain [21].

In conclusion, B cell-IgG-FcγR signaling drives the development of mechanical allodynia following CCI or NC, but not SNI or SNL. While secretion of autoreactive, pronociceptive IgG appears to be a generalized response to traumatic nerve injury, the contributions to neuropathic may be regulated by differential levels of the FcRγ subunit in the DRG and spinal cord. Our previous work implicated B cell and IgG signaling in the DRG from donors with neuropathic and other forms of chronic pain [21; 46]. The preclinical data presented here point to a need to develop strategies to identify those patients whose pain is functionally driven by FcγR signaling. Such strategies may guide potential use of immunotherapies targeting B cell-IgG-FcγR signaling to treat neuropathic pain.

## Conflict of interest statement

The authors declare that no conflict of interest exists.

## Acknowledgements

This work was funded by National Institutes of Health grant R01NS126252 (P.M.G. and C.J.H.) and in part by P30CA016672 (animal housing and care in the MD Anderson Research Animal Support Facility). The murine anti-CD20 monoclonal antibody and IgG2a isotype control antibody were generously gifted by Genentech under a material transfer agreement. We thank Microarray Phenotyping Core at UT Southwestern Medical Center (Dallas TX) for their help with Autoantigen array printing, processing and data generation. We thank Monoclonal Antibody Core at UT MD Anderson Cancer Center for their help with serum IgG isolation and purification.

## Author contributions

N.T.F., C.J.H. and P.M.G. conceived the study. N.T.F, K.F.W., C.J.H., and P.M.G. designed experiments. N.T.F., K.F.W., A.R.G., D.D., and Y.A.Z., collected the data. N.T.F., K.F.W., C.J.H., and P.M.G. drafted the paper. All authors contributed to data analysis and writing of the paper. Authors declare that they have no competing interests.

## Notes

### Competing Interest Statement

The authors have declared no competing interest.

