## Supplementary materials for "Autoreactive IgG levels and Fc receptor γ subunit upregulation drive mechanical allodynia after nerve constriction or crush injury"

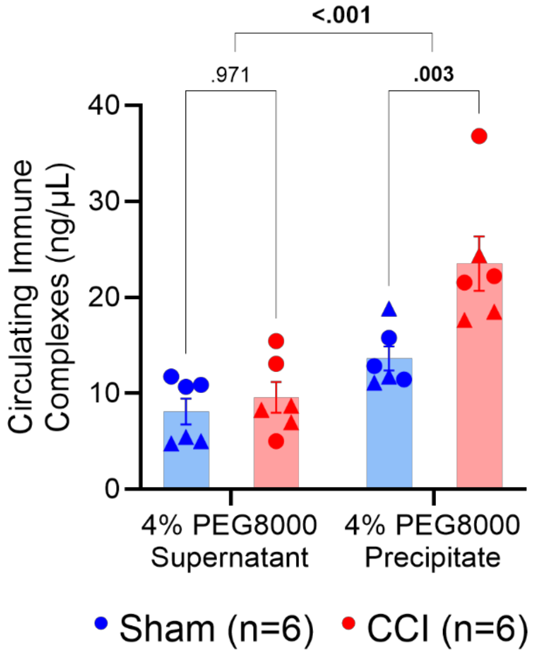


**Figure S1. CCI increases circulating IgG ICs.**

Quantification of circulating IgG ICs protein levels isolated from serum following 4% PEG8000 precipitation via ELISA. ELISA analyzed by two-way ANOVA and Dunnett's post hoc test; *p* values shown for comparisons CCI vs Sham and Precipitate vs Supernatant. *n* = 6/group (3 males and 3 females). For all data points, circles (●) represent male and triangles (▲) represent female mice.

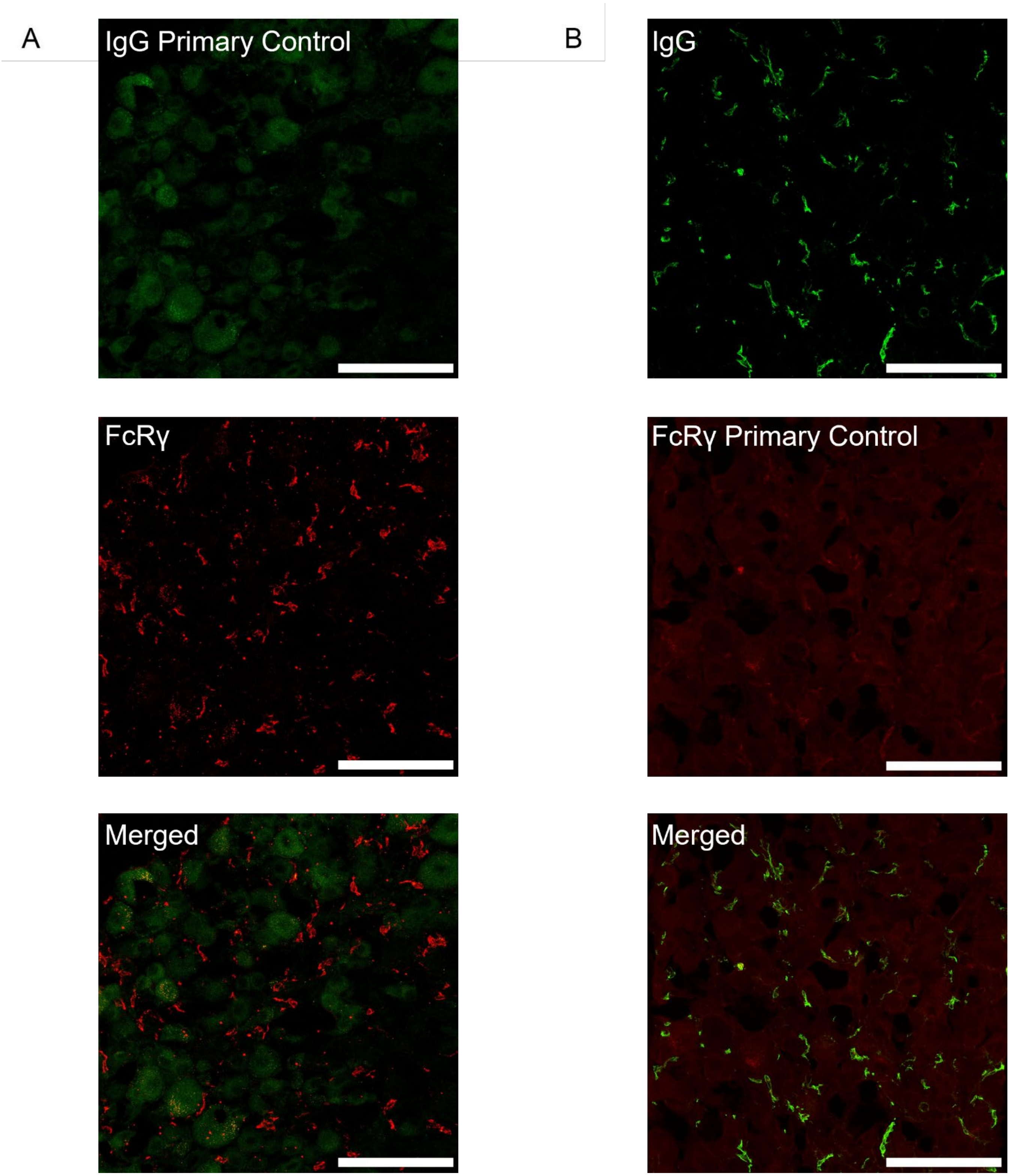


**Figure S2. IgG and FcRγ antibody controls.**

(**A**) IgG and (**B**) FcRγ antibody specificity was confirmed using primary antibody controls in DRG WT mice. Representative 40x fluorescent images (male, day 14 after surgery). Scale bars indicate 100μm.

**Table S1. List of antibodies and primers.**

| Antibody / Primer | Vendor | Catalog# | Dilution/Conc |
|---|---|---|---|
| Goat anti-mouse IgG, $F(ab')_2$ fragment specific | Jackson ImmunoResearch | 115-005-072 | 1:100 |
| Rabbit anti-Map2 | Abcam | ab32454 | 1:250 |
| Mouse anti-FcRγ | MBL Life sciences | M191-3 | 1:200 |
| Rabbit anti-NK.1 | Invitrogen | PA132229 | 1:100 |
| Donkey anti-goat IgG (H+L), Alexa Fluor 488 | Invitrogen | A-11055 | 1:500 |
| Donkey anti-rabbit IgG (H+L), Alexa Fluor 488 | Invitrogen | A-21206 | 1:500 |
| Donkey anti-mouse IgG (H+L), Alexa Fluor 594 | Invitrogen | A-21203 | 1:500 |
| Donkey anti-rabbit IgG (H+L), Alexa Fluor 594 | Invitrogen | A-21207 | 1:500 |
| DAPI (hydrochloride) | Sigma Aldrich | D9542 | 1:5000 |
| *Fcer1g* Forward Mouse: GTATTGTCCTTACCCTACTCTAC | Sigma Aldrich | KSPQ12012G | 500nM |
| *Fcer1g* Reverse Mouse: TGCTTCAGAGTCTCATATGTC | Sigma Aldrich | KSPQ12012G | 500nM |
| *Gapdh* Forward Mouse: GTTTGTGATGGGTGTGAACC | Invitrogen | A15609 | 500nM |
| *Gapdh* Reverse Mouse: TCTTCTGAGTGGCAGTGATG | Invitrogen | A15609 | 500nM |
